# Evidence for alternate stable States in Collapsing Ecological Networks

**DOI:** 10.1101/046045

**Authors:** Suresh Babu, Gitanjali Yadav

## Abstract

**Background:** There has been considerable interest and progress in our perception of organized complexity in recent years. Recurrent debates on the dynamics and stability of complex systems have enriched our understanding of these systems, but generalities in the relationship between structure and dynamics are hard to come by. Although traditionally an arena for theoreticians, much of this research has been invigorated by demonstration of the existence of alternate stable equilibria in real world ecosystems such as lakes, coral reefs, forests and grasslands.

**Results:** Linking up systems thinking with recent advances in our understanding of ecological networks opens up exciting possibilities. In an attempt to obtain general patterns of behaviour of complex systems, we have analyzed the response of eighty-six real world ecological networks to targeted extinctions, and the findings suggest that most networks are robust to loss of specialists until specific thresholds are reached in terms of geodesics. If the extinctions persist, a state change or ‘flip’ occurs and the structural properties are altered drastically, although the network does not collapse. Further, we find that as opposed to simpler networks, larger networks have several such alternate states that ensure their long-term persistence and that indeed complexity does endow resilience to such networks.

**Conclusions:** This is the first report of critical transitions in ecological networks and the implications of these findings for complex systems characterized by networks are likely to be profound with immediate significance in conservation biology, invasion biology and restoration ecology.

## BACKGROUND

Interest in ‘robust, yet fragile’ nature of complex systems transcends disciplinary domains of biology, engineering, sociology, economics, and ecology[1,2]. There is much to be gained by investigating the behavior of unique complex dynamical systems like ecosystems that are robust by virtue of their continued existence in evolutionary time[2]. The structural attributes shared by these systems could provide clues about their stability and robustness. Systems such as scale free ecological webs display an unexpected degree of tolerance or structural robustness to loss of specialists[3,4]. Studies on mutualistic networks have highlighted that modularity-one of the emergent properties of networks, endows robustness[5]. Be it fire prone savanna ecosystems, spread of infectious diseases or financial networks like the Fedwire, compartmentalization has been shown to render the much needed robustness to these systems[2]. The dynamics of large complex networks formed by interacting species impact the way biodiversity influences ecosystem functioning[6]. Understanding the behaviour of ecological networks is also central to understanding the response of biodiversity and ecosystems to perturbations.

Although there is adequate evidence to imply that structural and topological attributes of networks influence dynamics and function[7,8,9], the attributes of nodes and overall topological properties of networks that endow stability against perturbations are not sufficiently understood. The ‘targeted extinction’ approach for exploring the effects of node loss and associated co-extinctions has been well established over the last decade[3,10,11]. This approach involves simulations of random or ordered primary extinctions based on a given node property such as the number of links or ‘degree’. The response of the network in terms of resulting secondary extinctions or other network properties can be used to infer the significance of the node attribute being studied[4,12]. Extensive work on the robustness of ecological networks and attributes that enable species coexistence[5] and diversity[9] have revealed that these networks are highly robust to loss of specialists but are unable to withstand the targeted removal of generalists. This study was undertaken with the aim to understand how species persist in a collapsing mutualistic network following targeted species removals and associated coextinctions. The initial analysis was carried out using primary data on frugivory collected from Great Nicobar Island, India (GNIC) and the response and behavior of the network was explored in terms of established network attributes including species richness, secondary extinctions, nestedness, fragmentation and diameter. The analysis led to the discovery of alternate stable states that help sustain the integrity of the collapsing network. These alternate states are identifiable in terms of two attributes describing internal communication within the network, namely, its diameter (LDia) and number of diameters (NDia). A comparative analysis using extinction cascades across eighty-five ecological networks of varying nature was used to validate the observations made with GNIC, which revealed that these alternate states and flips were indeed pervasive across all the networks.

## RESULTS

The GNIC is a single large connected network having 812 interactions between 38 frugivores and 181 tree species (binary interaction matrix in Supplementary Data A1), making it one of the largest frugivory networks reported to date. Figure 1a represents an edge weighted, force directed layout of GNIC depicting highly asymmetric interactions, a characteristic path length of three, and a diameter of six. A radial layout of GNIC showed that species interact with nested subsets of partners. As expected of ecological networks, the nestedness of GNIC is high (NodF value 21.02) and its degree distribution showed best fit to a truncated power law distribution (Figure 1b and c). In comparison with other frugivory webs reported to date, GNIC has higher links per species (L/S), greater density, asymmetry and specialization, and a comparatively lower Connectance (L/S^2^).

**Figure 1.**
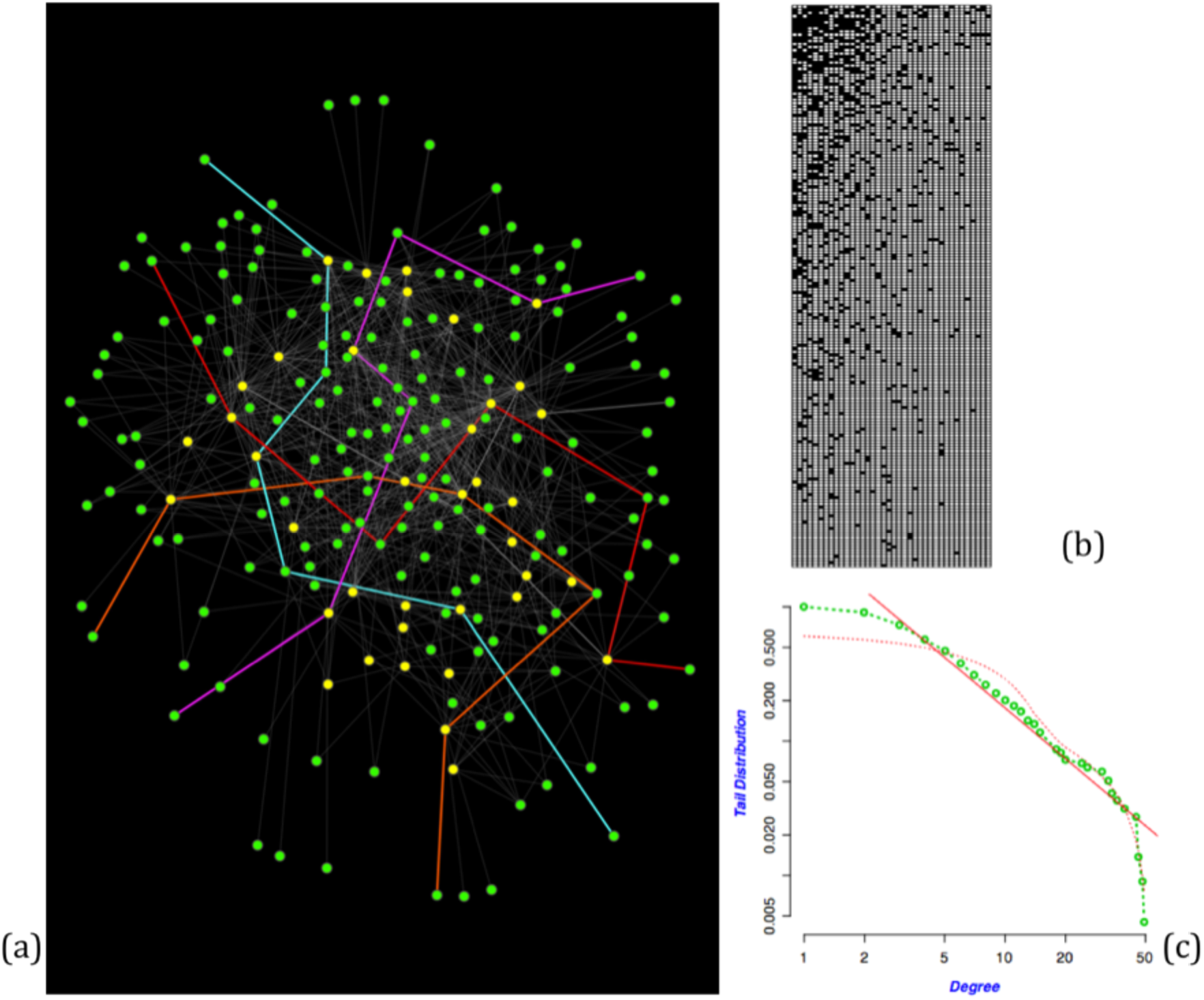
Illustration of the GNIC frugivory network from Great Nicobar Island, India: (a) Visual layout - circles represent species or nodes (green for plants; yellow for animals) and lines or edges between nodes represent the interactions between species. GNIC is a bipartite network with a diameter or LDia of 6. Four diameters are highlighted in this figure as thick paths of length 6, in blue, orange, red and pink. The unperturbed network contains 10065 such diameters or NDia, i.e independent shortest paths of length 6, between a given pair of nodes, (b) The nested pattern of GNIC matrix of interactions, and (c) Degree distribution of GNIC nodes, showing the best fit to truncated power law curve.

### Co-extinction Analysis

Degree based co-extinction simulations were carried out for GNIC, and the response was observed using different network level indices. As species removal is simulated from the most specialized (least-linked) to most-generalized (most-linked), along a specialist-generalist continuum, species richness decreases linearly and secondary extinctions do not occur until 61% primary extinctions. This can be seen in Figures 2a-b, which also depict the opposing ‘generalists-first’ extinction sequence, resulting in a sharp decrease in species richness accompanied by drastic secondary extinctions. This contrasting response was also observed in terms of the overall change in nestedness after perturbation; as specialists are removed, nestedness of the resulting networks tends to increase, while the removal of generalists triggers a rapid loss of nestedness in the corresponding reduced webs (Figure 2c). Notably, GNIC was able to sustain its unfragmented nature throughout the specialist-first extinction sequence, disintegrating only after removal of over 90% nodes; whereas, the reverse sequence (generalist first extinction) resulted in catastrophic network fragmentation into many disconnected sub-webs and complete collapse within the first 23% primary species removals (Figure 2d).

**Figure 2.**
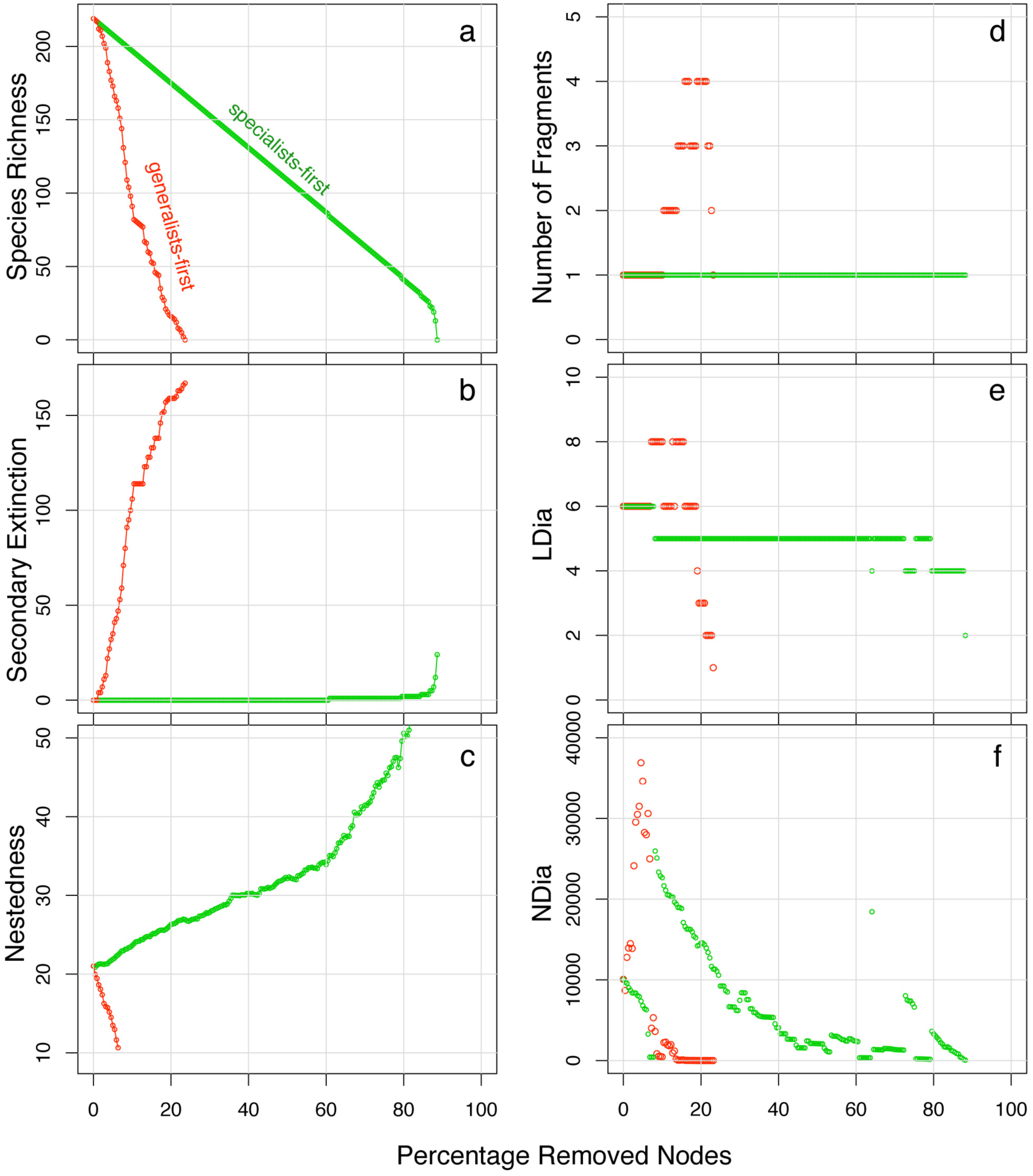
GNIC response to targeted extinction in terms of six distinct network attributes resulting from generalists-first (red) and specialists-first (green) extinction sequences are plotted against the fraction of nodes removed. These six attributes are (a) Species richness (b) Secondary Extinctions (c) Nestedness (d) Fragmentation (e) LDia and (f) NDia. Note the coupled variations between panels d-e-f at a given point on the X-axis. In the generalists-first scenario, when NDia is at its lowest, the reduced network fragments into disconnected subwebs (by 10% loss) followed by rapid collapse. However, in case of specialists-first, whenever the NDia reaches a minima, there occurs a compensatory reduction or ‘flip’ in LDia, causing an immediate recovery in NDia values, and the unfragmented nature of the network is preserved.

To explain the observed robustness of GNIC to loss of specialists, we carried out a detailed examination of the two contrasting breakdown scenarios, in terms of additional bipartite network attributes such as degree-distribution exponent gamma, density, asymmetry, connectance, generality, specialisation, C-score, V-ratio and various aspects of geodesics, including number, length and unique sets of shortest paths. Although for most of these properties, patterns were not discernable, two mutually independent attributes describing internal network communication (LDia and NDia) showed striking structural breaks (Figure 2e and f). Most prominently, these two attributes show a coordinated response when specialists are removed, revealing a characteristic pattern that is absent in case of generalists-first extinctions. As can be seen, in the generalists-first extinction sequence, LDia briefly increases followed by a steep decrease and rapid collapse, a trend that corresponds with NDia curves; loss of generalists leads to a brief but drastic increase in NDia (to over 36900 at just under 5% primary extinctions), after which it steeply drops (to 455 by 10% deletions). This low number of diameters (NDia) corresponds to a failure of internal communication and subsequently the network undergoes fragmentation. A comparison of the plots (Figure 2d and e) shows that the first instance of fragmentation in the collapsing network occurs at about 10% deletions, coinciding exactly with the lowest value of NDia. Further node deletions rapidly result in more fragments and the network collapses by 23% removals. In contrast, when specialists are removed first, LDia remains constant and NDia decreases steadily. By about 8% deletions, NDia reaches its lowest value of 436. However, the plot in Figure 2d shows that despite minimal internal communication, the single unit connected character of the collapsing network is preserved. Interestingly, the lowest value of NDia corresponds to a single unit reduction in the LDia, which in turn, results in a steep recovery of NDia values (from 436 to 25966). Subsequently this pattern repeats itself, i.e, NDia decreases at pace with loss of specialists till about 80% extinctions. At its lowest value, it drastically rises again - corresponding to a further unit reduction in LDia. Evidently, the coordinated response between NDia and LDia and the associated renewal of internal communication, makes it possible for the reduced network to make a stable transition to a new state and remain unfragmented, all through the specialist-first extinction sequence. Such a compensatory ‘flip’ response between two network attributes, specific to the specialist-first scenario, and absent in the generalist-first scenario, has not been reported before. Supplementary Data Sheet A3 provides details of this analysis, and the number of fragments, co-extinctions, LDia and NDia measured after every consecutive species deletion, both for the specialist-first and the generalist-first extinction cascades.

### Comparative Analysis of Ecological Networks

A comparative analysis of 85 additional ecological networks showed these patterns in LDia, NDia and fragmentations to be consistent and pervasive across all networks during the specialists-first breakdown scenario, and not limited to mutualistic webs only. As with GNIC the coordinated variation between NDia and LDia values endowed robustness to the perturbed networks and they persisted as single connected units during the attacks. In several cases, the different states were more pronounced than observed for GNIC. At least two and upto six flips were observed across the networks. In networks with low interaction density (< 1.35), the transition between states was not very clear. In cases where the initial network was disconnected, the specialist-first extinction sequence began with the removal of the smaller unit/s, and eventual persistence of the single largest unit. Figures 3a and b depict the synchronised behaviour observed in two of the 25 frugivory webs studied (codes SILV and JOR1, see Supplementary Data for details). Figure 3c shows the flips observed in MEMO, a pollination network with 299 interactions among 104 species. Figures 3d and e depict similar plots for an anemone-fish network (ANEM), and an ant-plant network (BLUT) respectively. Figures 3f-h depict the LDia-NDia plots for a host-parasite, plant-herbivore and a predator-prey network (LAKE, JEOM and MART) respectively.

**Figure 3.**
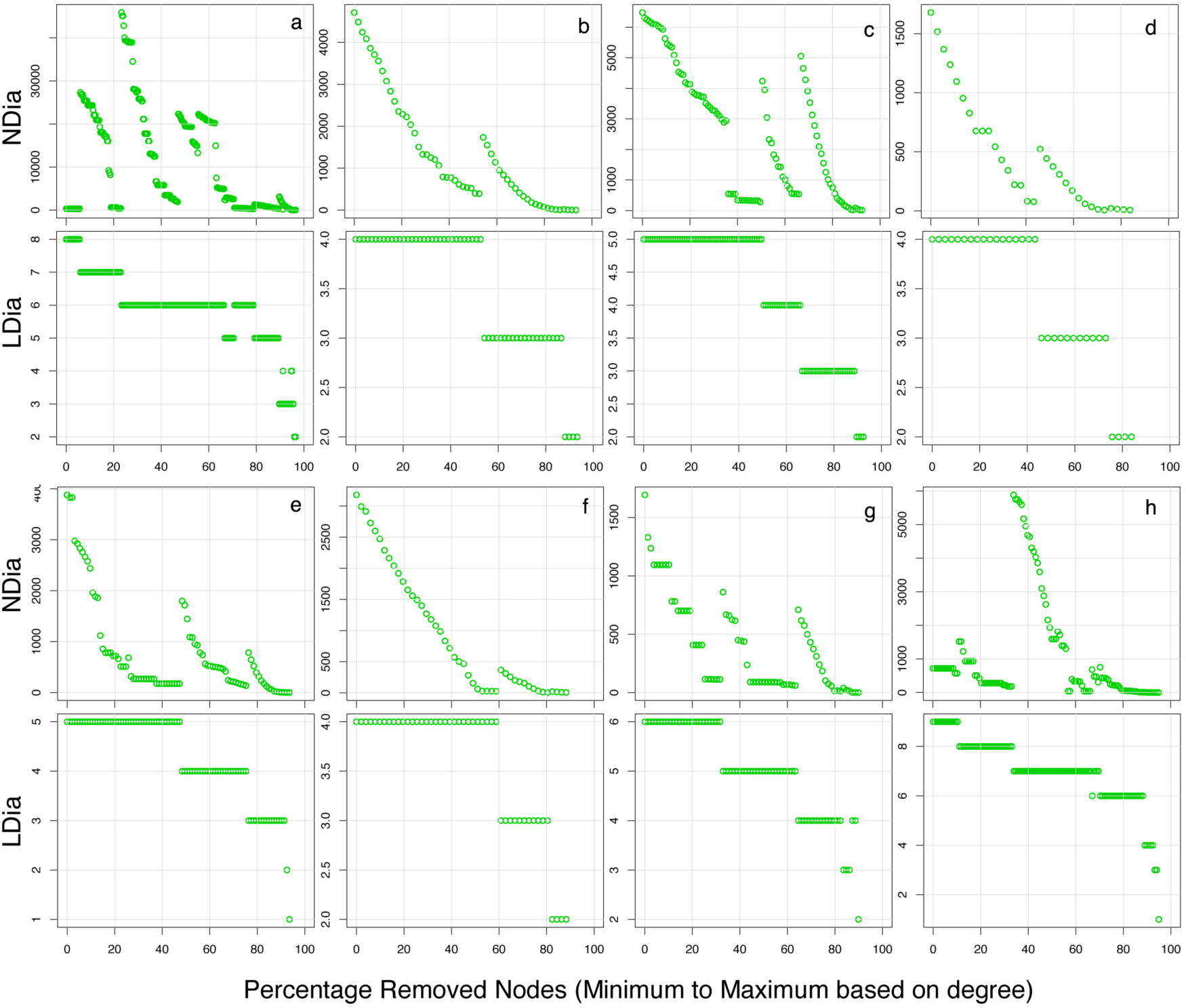
Variation between NDia and LDia in eight representative ecological networks following extinction cascades, resulting in flips and alternate stable states. Panels *a* and *b* depict two seed dispersal networks (SILV, JOR1) while panels *c* to *h* depict one each of pollinator (MEMO), Anemone-Fish (ANEM), Ant-Plant (BLUT), Host-parasite (LAKE), Plant-herbivore (JEOM) and Predator-prey (MART) network. For each network, the panels contain the corresponding NDia and LDia plots arranged vertically below each other. As in Figure 2, each plot depicts the change in the respective network attribute as a function of the fraction of nodes removed. Note that each drop in LDia corresponds to a surge in the NDia for a given web. The length/duration of each stable state can be measured as the fraction of nodes removed while the LDia remains constant. The number of states can be measured as the number of stable flips in LDia. Details for each network are provided in Supplementary Data A2.

For all networks, ‘resistance to flip’ was estimated in terms of percentage of primary species extinctions after which the first flip was observed. Therefore ‘resistance to flip’ is higher if a large number of species deletions are required before the flip is observed, and lower if fewer node deletions cause the state change. Figure 4a shows the relationship between the ‘resistance to flip’ and initial LDia across the 86 ecological networks. Compact networks with small diameters require over 70% primary extinctions for a flip in their state (Figure 4a). Networks with initial diameters of 6 or more require a much smaller proportion of primary extinctions to switch to lower diameters. A linear regression of ‘resistance to flip’ on initial LDia indicates a significant negative relationship (Adjusted R^2^ = 0.3598, β = −7.455, df=84, p-value: 6.288e-10). However a Loess plot of the same indicates a curvilinear rather than a linear relationship between the variables, when examined at different spans (Figure 4b). The data was found to meet the assumptions of homoscedasticity and normality of errors (Supplementary Figure S1). It was also observed that a positive, nearly linear relationship exists between the likely number of flips and the initial diameter of the network (Linear model; Adjusted R^2^ = 0.5981, β = 0.559, df = 84, p-value: < 2.2e-16; Supplementary Figure S2). Results of the entire analysis of 86 ecological networks are presented in Supplementary Data A.

**Figure 4.**
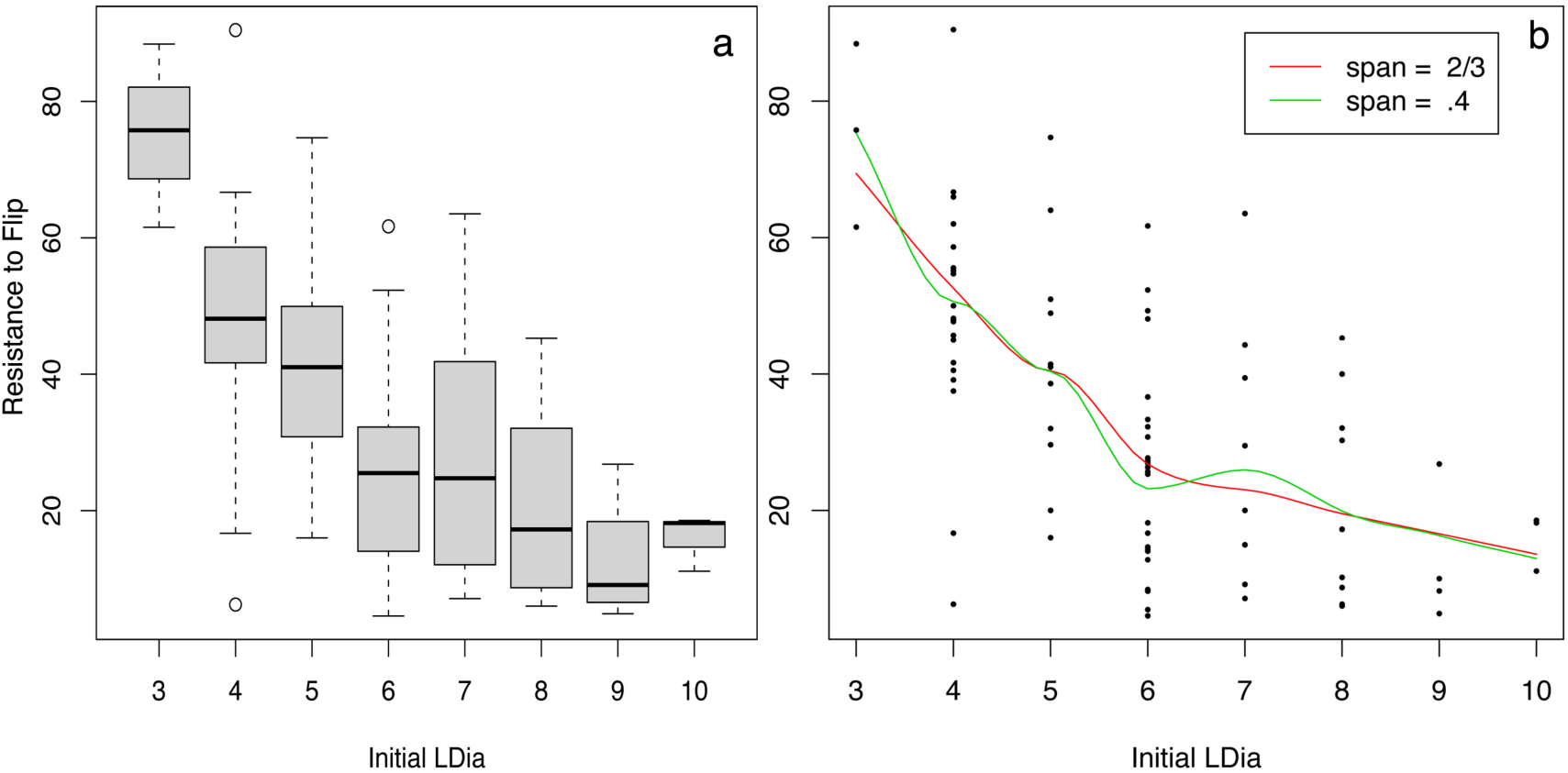
Influence of initial LDia of a network on its ‘resistance to flip’ or the primary extinctions required for a flip. The box plot provides a summary of the observations for 86 networks analysed in this study (a). Networks with smaller initial LDia show resistance to flip compared to networks with larger LDia as more primary extinctions are needed for a state change. Loess of the same (b) suggests a curvilinear rather than linear nature of the relationship. The larger networks have lower resistance to flip indicating that fewer primary extinctions can cause a state change in their communication levels.

## DISCUSSION

Ecological networks are well known to be robust to removal of specialists, but reveal an intrinsic structural fragility in response to targeted removal of generalists, eventually resulting in fragmentation into many small sub-webs. This behavior is not shared with random networks, which are equally fragile to random or selective node removals[13]. We began this work with GNIC frugivory data, a new network showing characteristic features similar to known ecological webs, to investigate the contrasting ability of mutualistic networks to withstand attacks on specialists as against generalists, the former known to be a more realistic extinction threat. As expected, distinct responses were observed. Generalist first extinction cascade of GNIC caused the species richness to plummet due to steep rise in coextinctions, whereas the specialist first cascade shows a linear decrease in species richness as it does not involve the loss of associated species. Nestedness, one of the most significant and widely observed non-random pattern in networks of ecological interactions, is known to greatly affect the robustness of mutualitsic networks[14]. The variation in the nested structure of the reduced webs in the two contrasting scenarios revealed that as specialists are removed, nestedness of the reduced network increases, while the removal of generalists resulted in a steep decrease in nestedness, supporting the notion that nestedness provides alternate routes for system responses after perturbations such as species extinctions or link removals. Nestedness being a measure of robustness[9,14] this indicates that extinctions of specialists improve the robustness of the reduced networks. Of the several structural attributes analysed to understand this ability, the most striking patterns emerged from the behaviour of the geodesics as the network was subjected to systematic extinctions.

### Emergent properties of collapsing networks

The coextinction cascades simulated with GNIC showed that with specialists being removed first, the graph does not fragment unless substantial primary extinctions have occurred. Our initial findings suggested that in the specialist-first scenario, network attributes ‘re-wire’ to make the reduced network more compact, thereby maintaining optimal communication between the remaining nodes, keeping the network unfragmented. These topological re-adjustments are characteristic and were identified in terms the network diameter, which represents the longest geodesic of the network. Although the diameter has been relatively less studied in mutualistic webs, it is a well established measure of topological robustness of several complex communication systems, ranging from cells to social, civilian networks and the Internet[10,15]. For a given network, a low diameter is considered advantageous as it can contribute to greater interconnectedness, shorter communication paths and lower load on links, or edges. We examined two aspects of the diameter: (a) its length or ‘LDia’, and (b) the number of diameters or ‘NDia’, and discovered a striking synchrony between LDia and NDia in the specialist-first extinction scenario, presumably an internal compensation that endows the perturbed network with the ability to avoid fragmentation. Every instance of a very low NDia value was found to coincide exactly with a corresponding reduction in the LDia value, leading to a reversal of the decreasing NDia trend. A sufficient number of NDia in the collapsing network presumably enable it to maintain communication between remaining nodes, which remain connected despite the sustained perturbations. This coupling was not observed in the generalist-first extinction sequence, where the perturbed network, unable to recover after an excessive decrease in its NDia, undergoes multiple fragmentations.

We have attempted to explain the contrasting responses in the two breakdown scenarios using a schematic in Supplementary Figure S3. As shown in this Figure, the generalist-first extinction sequence begins with a loss of crucial hubs through which the original diameter-paths were running, and this deletion leads to the selection of longer routes, causing the LDia to increase at first. Further loss of hubs then causes the network undergo fragmentation into clusters of nodes, whose sizes (and corresponding LDia) decrease rapidly with continued node removal, until total collapse. In contrast, when the specialists are removed first, the nodes break away one by one from the periphery, rather than as clusters, so that a large number of alternate paths of length LDia remain available. As a result, the NDia slowly decreases while the LDia remains constant. As NDia reaches its lowest value, the LDia flips and becomes smaller. This reduction in LDia causes an abrupt rise in NDia is periodic and enables the network to avoid fragmentation when the NDia reaches extremely low values. We ascribe this rise in NDia to the combined number of (a) pre-existing paths at the new (reduced) LDia, and (b) the node loss driven shorter paths created from the previous (larger) LDia. This LDia - NDia synchrony repeats itself all along the breakdown scenario although on a progressively smaller scale. Consequently, the network displays high topological stability throughout the extinction cascade by remaining un-fragmented.

The iterating pattern of gradual decrease in NDia till a threshold of extinctions is reached, followed by a sudden transition to a new high value at a lower LDia, resembles the behaviour of ecosystems that can exist in multiple states characterized by unique sets of conditions[16,17,18]. The theory of alternate stable states suggests that the discrete states are separated by thresholds and the system remains in one state unless perturbation is large enough to tip it over to the next state[16,19]. It has been suggested that gradual changes are more like the rule and critical transitions are an exception, which demand special attention[20]. In case of GNIC, the network retains its integrity by flipping between alternate levels of communication and complexity expressed in terms of LDia. However for the reduced network, the increased NDia now endows the system with high resilience, as the threshold required for the next flip or shift in LDia requires over 60% primary extinctions. The state with the widest stability basin, characterized by the maximum range of NDia at a given LDia, provides much of the robustness of the network. Figure 5 depicts this in a schematic representation. Hysteresis or path dependency, characteristic of alternate equilibria, becomes evident once a flip in LDia has occurred. If the lost node were to be returned to the network at this stage, it may not bring the system back to the previous state. Rather, it would lead to an increase in the NDia within the current state, i.e at the new value of LDia. This is likely because of the increasingly nested pattern of the reduced networks (Figure 2c) a new species is likely to preferentially attach to the ‘hub’ nodes or generalists[6,21,22]. Attachment to a hub node does not lead to an increase in LDia; it can only result in additional alternate paths or NDia. As a result, the system will not return to its previous state just by a simple reversal of extinction, or re-introduction of lost species. This observation may have wide implications in the area of restoration ecology and invasion biology, as we discuss later.

**Figure 5.**
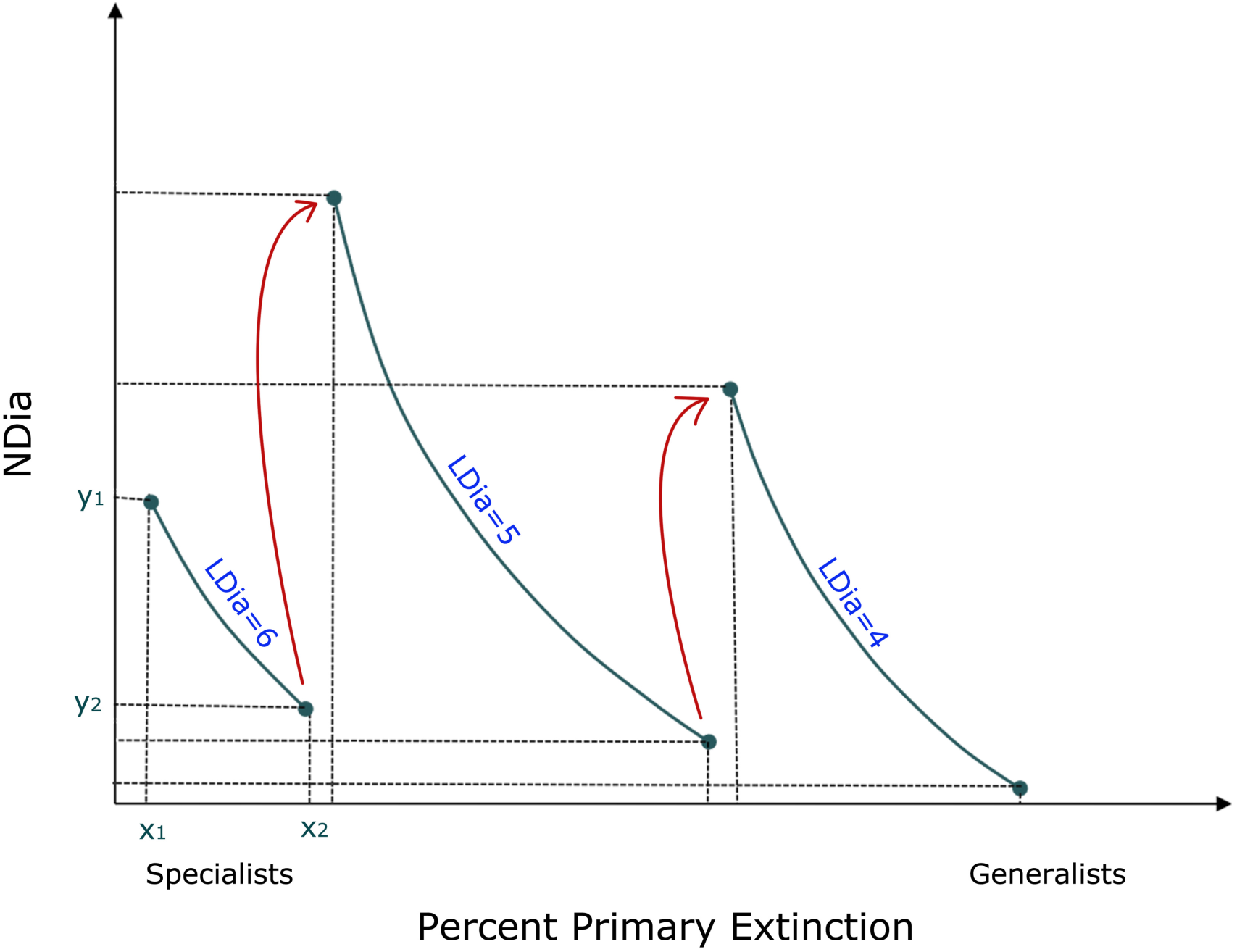
A schematic framework explaining the alternate states following a specialist first extinction cascade in ecological networks. This is a schematic representation using GNIC as the baseline. The X-axis corresponds to percentage primary extinctions with specialists being removed first along a specialist-generalist continuum. The Y-axis corresponds to the number of diameters or NDia values of the network. Each state is depicted as a grey arc. For instance, x1 corresponds to the unperturbed status of the network (percent primary extinctions=0) and x2 corresponds to the threshold value of percent primary extinction beyond which a flip (shown by bold arrows) occurs to the next state. The range of NDia values that correspond to the primary extinctions (y1 to y2) characterize the resilience of the network as it maintains its state (LDia=6) until it reaches a threshold of extinctions at x2, beyond which a state change to lower diameter (LDia=5) occurs. This shift is path dependent since addition of a node at this stage is unlikely to bring about a reversal of the state (see discussion), resulting in hysteresis as state changes occur repeatedly in the cascade.

The following generalizations emerge from our observations on GNIC: (a) total number of alternate paths or diameters (NDia) decrease with loss of specialists, (b) for a given network, reduction in the diameter (LDia) increases the NDia, (c) the LDia reduction occurs only at, or beyond, a critical loss of specialists, (d) the network precludes fragmentation, with the loss of specialists across the entire cascade and (e) for a given network, there may be several alternate stable states that can spring surprises against slow moving perturbations which can be masked by internal adjustments of the network.

The generality of these observed patterns in LDia-NDia was established by a comparative analysis across 85 additional ecological networks including mutualisms as well as antagonistic. webs. Our results show that the collapsing network sustains its connected or un-fragmented nature during the loss of specialists by internal structural readjustments in terms of LDia and NDia, which is not evident during the loss of generalists, thereby leading to immediate collapse. We also find that initial network size corresponds to the number of flips observed. Larger networks are likely to have more number of alternative stable states to cope with uncertainties in evolutionary time. For example, a small network like the anemone-fish network has only 36 species and an unperturbed LDia of 4, resulting in only one alternate stable state which may restrict its ability to withstand perturbations (Figure 3d). Larger networks like the Brazilian Amazon (code SILV) and GNIC have several possible alternate stable states and are more likely to persist under long periods of adversity. Smaller perturbations tend to flip larger, more complex networks to alternate states (Figure 4a and b) and since they have several such possible states, the network architecture endows resilience to such networks. The smaller, less complex networks do not show any state changes under small perturbations indicating resistance. However, since smaller networks also have very few possible alternate states, they are low on resilience. The width of stability basins and the number of possible stable states that accompanies the loss of specialists progressively shrinks, as the network size reduces, thus affecting its overall resilience. Therefore there may be an evolutionary advantage in making ever larger webs of interactions that facilitate long-term persistence of species rich communities, a finding that complements a recent study[9] as to how mutualistic communities can enhance co-existence of species.

### Implications of alternate states in ecological networks

Co-extinctions are now recognized as a major driver of global biodiversity loss, along with habitat destruction, species invasion and overkill [23,24]. Since more than half of all known species and a large proportion of unnamed ones are involved in host specific relationships in atleast some stage of their life, specialists face a greater risk due to secondary extinctions[25,26]. There is added relevance of re-examining the threats of extinction knowing that interacting species may exist in alternate states. Besides broad implications on our understanding of bipartite networks in general, our findings have significance in conservation biology, invasion biology, and restoration ecology. Based on the present positioning of an interaction network along an extinction cascade, it may be possible to predict the proximity of the system to a catastrophic change and model real time stability indicators of networks. In addition to the dynamics associated with ‘critical slowing down[27], this may be an alternate approach to predict the likelihood and proximity of a system to regime flips. Conservation programmes could benefit from directly identifying the most threatened systems, requiring immediate attention or prioritization.

High nestedness and NDia of the reduced networks can make the system receptive to invasive species because of the benefits associated with joining a well-connected network. Invasive species have been shown to be able to take advantage of existing mutualistic networks in invaded habitats[28] and the state of the native network may explain its invasibility. A native seed dispersal network in which most of the specialists have been removed is more vulnerable to invasion as the invader would be able to associate with a generalist without much competition from other specialists. This provides the invader with enhanced connectivity, which has a vital role in its persistence and spread. Coupled with experimental evidences[22] our findings pave the way for developing a network approach to invasion biology.

Hysteresis in collapsing networks as implied in the theory of alternate stable states has for instance profound implications in restoration ecology. It may be possible to explain why certain restoration programmes do not follow expected trajectories and one may aspire to find system specific predictors of thresholds of recovery. Path dependency of collapsing networks informs us about the near impossibility of reconstructing highly degraded ecosystems.

Another functional consequence of such a pervasive phenomenon would be on the ability of networks to transmit or ‘percolate’ perturbations across the network. Coupled oscillations are likely to travel far and wide across the network much more effectively as LDia decreases. In real ecosystems, this could have major implications. A drastic reduction in the abundance of a particular species owing to hunting or disease would impact the network much faster in unpredictable ways. This would again imply an increased uncertainty over the behaviour of such networks. The flips in diameter following loss of specialists could well upset the functional advantages of scale free networks[10,13]. The outward robustness of scale free networks to loss of specialists could mask the enhanced ability of the network to transmit perturbations.

## CONCLUSIONS

Our results provide empirical evidence for the direct link that exists between topological heterogeneity and system dynamics. We show by means of detailed analysis of eighty-six ecological networks of varying nature that the networks can exist in alternate diameters and levels of communication. The outwards stability and unfragmented nature of these networks against perturbations often mask the internal re-wiring that progressively reduces their resilience resulting in sudden flips or transitions to lower levels of communication. This study shows that the continuous loss of specialists leads to significant loss of resilience for the networks, which is irreversible - something impossible to demonstrate experimentally. On one hand these findings hint at an evolutionary advantage in building ever-larger interaction networks (moving to higher levels of robustness), and on the other hand also highlights the inability of heavily damaged networks to respond to restoration in tangible amounts of time. The increased likelihood of an invasive species attaching to generalists in an impoverished native network partly explains its success in invaded ecosystems. The robustness of scale free networks could disguise enhanced percolation of disturbances across the network. This study establishes a prevailing pattern across known complex ecological networks and open ups possibilities for empirically driven dynamical modelling of these networks. We expect our findings to be the starting point for an array of investigations into the importance of alternate states in ecological networks in particular and other kinds of networks in general.

### MATERIALS & METHODS

Primary data in the form of direct observations of foraging by vertebrates on fruits was collected from the tropical rainforests of Great Nicobar Island (spread from 6°45’ to 7°15’N and 93°38’ to 93°55’E, spanning a total area of about 1045km^2^), the southernmost Island in the Andaman & Nicobar archipelago, India, spanning a period of seven years with field work being conducted on fifty nine transects, each 500m long, in various regions of the island from December 1999 to November 2006. This study was undertaken as part of a larger initiative by the Ministry of Environment and Forests, Government of India, under the Man and Biosphere (MAB) Programme on Great Nicobar Biosphere Reserve, India. Direct observations of instances of foraging by vertebrates on fruits were recorded as an interaction matrix consisting of 181 plant species and 38 frugivores (33 birds and 5 mammals). Plant and frugivore species were identified and the interaction data obtained was compiled for the entire island. Data is presented as a binary interaction matrix (Supplementary Data A1). Preliminary analysis and visualization of network architecture was done using Cytoscape [29] version 2.6.2.

### Co-extinction Analysis

We simulated primary species loss by carrying out cascades of directed species removals or extinctions, based upon degree (the number of links), and compared the stability of the resulting reduced networks to random extinction cascades, following Memmott *et al*[3]. Upon removal of a species, those species that are left without any interaction are assumed to undergo co-extinction. The network remaining after each subsequent removal is assessed for robustness and stability. For each network, extinction sequences included both specialist-first (i.e least-linked to most-linked species) and generalist-first (most-linked to least linked species) cascades. Random removals were analysed after averaging from 300 replicates.

The network remaining after every primary extinction and subsequent co-extinction was analysed for its stability and robustness by measuring various network attributes commonly used to summarise and describe different patterns in ecological webs, such as degree, species richness, secondary extinction, fragmentation, lost interactions, degree-distribution-gamma values, axes, length and number of diameters etc. These indices were calculated using in-house fortran scripts and R CRAN packages IGRAPH[30] version 0.5, SNA[31] version 1.5, and BIPARTITE[32] version 0.91. Detailed description of each of the indices can be found within the respective package manuals. We examined the exponential, power law and truncated power law models to cumulative distributions for each network. Nestedness was calculated using the recently proposed nestedness metric NodF[33] using the ANINHADO program[34]. To assess the significance of nestedness values, the observed NodF was compared with benchmarks provided by three different null models. For each network, a population of n = 300 random networks was generated for each null model. As a statistic indicating significance, we estimated the probability, p, that a randomization was equally or more nested than the real matrix. Only the significant NodF values were used for further analysis. Comparison of nestedness across reduced networks was done without normalizing these values for variation in species richness or number of interactions, since each reduced network is essentially a subset of the original unperturbed network. The shortest paths (also called geodesics) were calculated by using breadth-first search in the graph. The diameter (LDia) of a graph is defined as the length of the longest geodesic. The number of diameters (NDia) was calculated as the sum of all diameters between every pair of nodes separated by a distance equivalent to the diameter (LDia). A unix program was designed to automate the entire analysis. This code takes a given binary network as input, simulates different co-extinction sequences and evaluates the sub-network remaining after every subsequent species removal, for its stability and robustness, and then extracts the attributes required for detection of regime flips or alternate stable states. For each reduced network, it creates a list of extinct and co-extinct species and calculates seven network level indices, namely species richness, secondary extinction, lost and remaining interactions, number of fragments, LDia and NDia and compares these indices across and between the different extinction sequences, and finally plots the results into vector format files. The source code is currently being developed as an open source web server facility.

In addition to GNIC, data records were obtained from a set of 85 ecological networks using previously published reports as well as the Interaction Web Database repository at the National Centre for Ecological Analysis and Synthesis (NCEAS) website (http://www.nceas.ucsb.edu/interactionweb). These 85 webs include one Anemone Fish network, four plant-herbivore, four ant-plant, seven host-parasite, one Predator-prey, 25 Seed dispersal or Frugivory networks and 43 Pollination networks. Each network was analysed using the code described above and subsequently examined for the occurrence of regime flips, as they appeared on plots of NDia and LDia with primary extinctions. Statistical analyses on the results across networks were carried out in R (V 2.11.0). Supplementary Table S1 contains a brief description of these networks and details with references are in Supplementary Data Sheet A2.

## LIST OF ABBREVIATIONS

### ACKNOWLEDGEMENTS

S.B acknowledges the Man & Biosphere (MAB) Programme of the Ministry of Environment and Forests, Govt of India, for support. G.Y thanks Director, NIPGR for support.

### FINANICAL DISCLOSURE

S.B acknowledges the Man & Biosphere (MAB) Programme of the Ministry of Environment and Forests, Govt of India, for financial and logistic support. G.Y thanks the Dept of Biotechnology (DBT), Govt. of India for funding. S.B acknowledges the research fellowship of Council for Scientific & Industrial Research (CSIR) in the initial years of this project.

### AUTHOR CONTRIBUTIONS

S.B designed the study and compiled the primary data used in the analysis. G.Y performed the computational simulations and wrote the co-extinction code used in this study. S.B and G.Y jointly analyzed the results and wrote the first version of the manuscript, both authors contributed to the final draft.

## SUPPLEMENTARY DATA FILES

**1. Supplementary Data A (EXCEL FORMAT FILE)**

This file contains Supplementary Data with sheets (A1) The Binary interaction data for GNIC frugivory network reported in this paper (A2) listing of all 86 datasets and their references (A3) detailed results of the co-extinction analysis on GNIC frugivory network, (A4) Summary results of the entire analysis on 86 networks.

**2. Supplementary Data B (PDF FORMAT FILE)**

This file contains Supplementary Figures S1, S2, S3, Supplementary Table S1.

### AUTHOR CONTRIBUTIONS

S.B designed the study and compiled the primary data used in the analysis. G.Y performed the computational simulations and wrote the co-extinction code used in this study. S.B and G.Y jointly analyzed the results and wrote the first version of the manuscript, and all authors contributed to the final draft.

